# Triggered reversible disassembly of an engineered protein nanocage

**DOI:** 10.1101/2021.04.19.440480

**Authors:** Jesse A. Jones, Ajitha S. Cristie-David, Michael P. Andreas, Tobias W. Giessen

## Abstract

Protein nanocages play crucial roles in sub-cellular compartmentalization and spatial control in all domains of life and have been used as biomolecular tools for applications in biocatalysis, drug delivery, and bionanotechnology. The ability to control their assembly state under physiological conditions would further expand their practical utility. To gain such control, we introduced a peptide capable of triggering conformational change at a key structural position in the largest known encapsulin nanocompartment. We report the structure of the resulting engineered nanocage and demonstrate its ability to on-demand disassemble and reassemble under physiological conditions. We demonstrate its capacity for *in vivo* encapsulation of proteins of choice while also demonstrating *in vitro* cargo loading capabilities. Our results represent a functionally robust addition to the nanocage toolbox and a novel approach for controlling protein nanocage disassembly and reassembly under mild conditions.

## Introduction

Intracellular compartmentalization is an effective strategy employed by all organisms to regulate metabolism and achieve spatial control.^1,2^ One widespread compartmentalization approach is the use of protein nanocages. They can accumulate and store labile compounds, sequester toxic or volatile reaction intermediates, and prevent undesired side reactions of encapsulated enzymes.^1,2^ Efforts have been undertaken to engineer protein nanocages like ferritins, lumazine synthase, and virus-like particles for various biomedical and industrial applications,^3–5^ but few have focused on engineering input-responsive nanostructures capable of triggered assembly or disassembly.^6,7^ Such controllable structures would expand the potential application range of engineered nanocages to include programmable delivery of encapsulated payloads and rationally timed substrate-product release and intermixing, to name only a few examples. Encapsulin nanocompartments have recently emerged as a particularly versatile bioengineering tool, resulting in their application as bionanoreactors, targeted delivery systems, and nano- and biomaterials production platforms.^8–11^

Encapsulins are icosahedral protein nanocages found in bacteria and archaea with triangulation numbers of T=1 (24 nm), T=3 (32 nm) or T=4 (42 nm) containing sub-nanometer pores at the symmetry axes.^12^ They self-assemble from a single HK97-fold capsid protein into 60mer (T=1), 180mer (T=3) or 240mer (T=4) protein cages and are involved in oxidative stress resistance,^13–16^ iron mineralization and storage,^17,18^ and sulfur metabolism.^19^ Their defining feature is the ability to encapsulate dedicated cargo proteins via short C-terminal targeting peptides (TPs) found in cargo proteins which specifically interact with the interior of the protein shell during self-assembly.^16,20,21^ This native feature has been reliably coopted for the facile encapsulation of non-native proteins through TP-fusions.^22^

Once assembled, encapsulins exhibit notable robustness and stability.^23,24^ While often a desirable characteristic, this also precludes their easy disassembly under physiological conditions, a key feature for responsive delivery systems, nanoreactors, and biomaterials. In particular, encapsulins' inherent stability prevents efficient release of molecules synthesized in their interior, cargo enzyme “hot-swapping” for sequential packaging, or triggered cargo release for drug delivery applications.

Here we develop an engineered protein nanocage based on a bacterial encapsulin that exhibits triggered reversible disassembly under physiological conditions while also maintaining cargo loading capabilities.

## Results and Discussion

### Protein cage selection and design of the disassembly trigger

The T=4 *Quasibacillus thermotolerans* encapsulin (QtEnc) was chosen as an engineering scaffold. QtEnc is the largest bacterial encapsulin known to date and is comprised of a thermostable, non-covalent chainmail formed from a single self-assembling protomer. Additionally, QtEnc is easily overexpressed and purified from *Escherichia coli* in an empty or cargo-loaded state.^18^ QtEnc was analyzed for engineerable structural features important for protein cage assembly that might also be tolerant to mutation, and would not interfere with cargo loading. We chose to focus on the elongated loop (E-loop) region of the encapsulin protein which makes critical intra- and inter-capsomer contacts and influences overall shell topology **(Figure 1A)**.^18^ The E-loop is also located away from the N-terminal helix important for cargo loading.^25^ Therefore, the E-loop was selected as the insertion site for the disassembly trigger.

**Figure 1.**
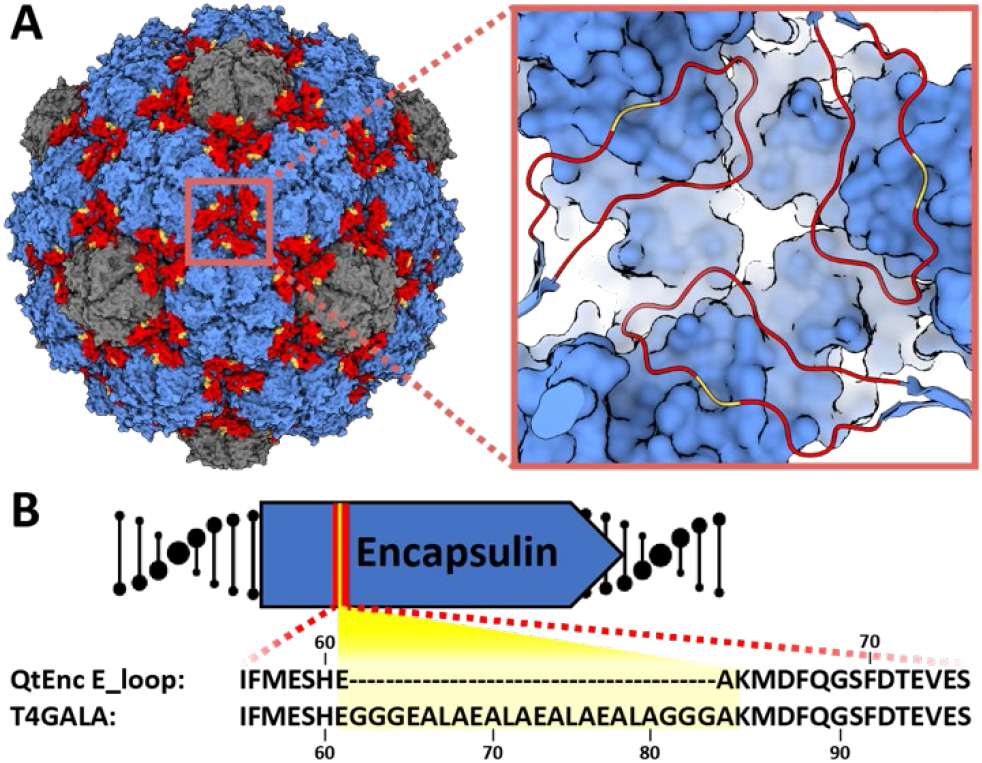
Design of the engineered protein nanocage. A) Surface view of the native *Quasibacillus thermotolerans* T4 encapsulin (QtEnc, PDB 6NJ8), highlighting hexameric (blue) and pentameric (gray) facets, and E-loops (red) along with the GALA peptide insertion site (yellow). Inset: zoomed-in view of the three-fold symmetry axis and insertion site. B) E-loop (red) sequence of QtEnc and T4GALA highlighting the GALA insertion (yellow).

The GALA peptide has been shown to demonstrate an inducible coil-to-helix conformational change upon acidification^6,26^ and was chosen as a disassembly trigger. A 16-residue GALA peptide flanked by triple glycine linkers was inserted between QtEnc residues Glu61 and Ala62 yielding the engineered nanocage T4GALA **(Figure 1B; Figure S1, TableS1)**. We hypothesized that under neutral and basic conditions, the GALA peptide random coil would not disturb E-loop conformation or shell assembly. Upon acidification, the GALA coil would be expected to adopt a helical conformation and introduce enough torsional strain to disrupt critical E-loop contacts, thereby perturbing structural integrity enough to induce disassembly of the protein cage. A reversion of the GALA helix back to its relaxed random coil state under less acidic conditions would be expected to allow reassembly of the encapsulin cage.

### Assembly, disassembly, and reassembly of T4GALA

To characterize the engineered nanocage, C-terminally His-tagged T4GALA was expressed and purified using Ni-NTA resin and found to still assemble via transmission electron microscopy (TEM) analysis (**Figure 2A**). Native polyacrylamide gel electrophoresis (PAGE) studies were then conducted to analyze the effects of pH, salt, and buffer on the engineered protein cage (**Figure S2**). T4GALA exhibited a tendency for disassembly at low pH, with near-complete disassembly achieved at pH 6.0. An unexpected dependence of T4GALA structural integrity on buffer identity was also observed. Specifically, disassembly at physiological pH was favored in the presence of Tris buffer (pH 7.5) while Bis-tris propane was found to significantly stabilize T4GALA under similar pH conditions.

**Figure 2.**
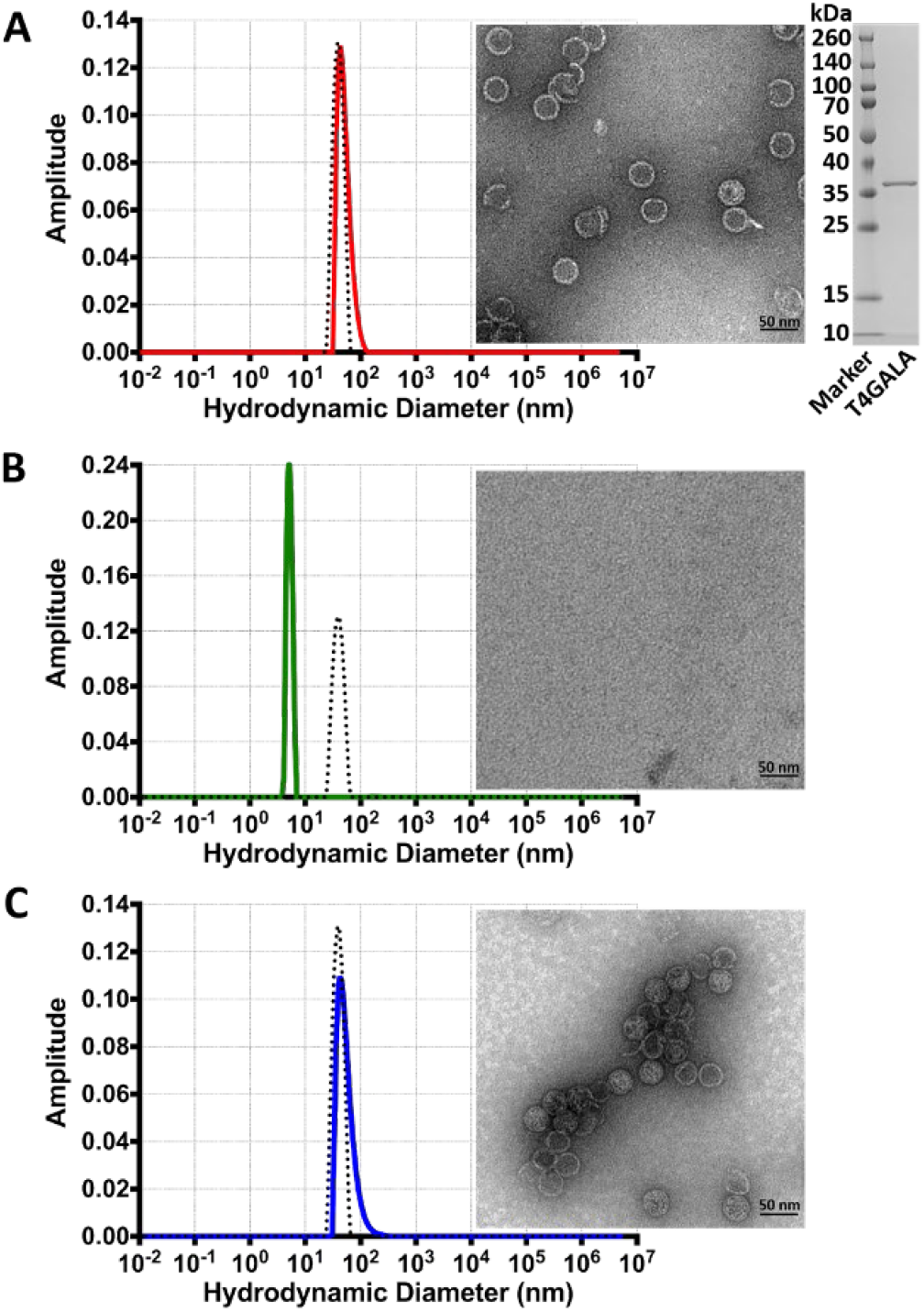
Assembly, disassembly, and reassembly of the T4GALA protein cage. A) Dynamic light scattering analysis (left) of assembled T4GALA (red) compared to native QtEnc (black dashed) with assembled T4GALA verified via TEM (right). SDS-PAGE analysis of purified T4GALA (far right). B) DLS analysis (left) of disassembled T4GALA after centrifugation (green) with QtEnc reference (black dashed) and disassembled T4GALA TEM analysis (right). C) DLS analysis (left) of reassembled T4GALA (blue) with QtEnc reference (black dashed) and reassembled T4GALA TEM analysis (right). Scale bars: 50 nm.

Size exclusion chromatography (SEC) showed that the elevated imidazole concentrations used for Ni-NTA elution helped maintain T4GALA in an assembled state even in Tris buffer. Imidazole was added to SEC buffers for all subsequent purifications (**Figure S3**). As such, T4GALA is easily overexpressed in *E. coli* and purified in the assembled state via a simple two-step protocol.

As concern existed regarding the potential for prolonged exposure to Tris buffer and unfavorable pH conditions during native PAGE analysis, assembly states were verified and characterized by a more reliable combination of dynamic light scattering (DLS) analysis and TEM **(Figure 2, Figure S4)**. A streamlined protocol was developed to purify T4GALA via standard Ni-NTA conditions, disassembly in imidazole-free Tris buffer, and reassembly in Bis-tris propane, all under physiological pH conditions. Overall, assembled T4GALA proved to be similar to native QtEnc in size (QtEnc Z-average diameter 47.2 nm, peak diameter 43.4 nm; T4GALA Z-average diameter 62.2 nm, peak diameter 56.39 nm) and monodisperse (**Figure 2A**), with the slight increase in average diameter by DLS possibly due to the additional disordered insert and potential small lipophilic aggregates. After brief centrifugation, the disassembled sample appears monodisperse with a diameter of ~6 nm (Z-average diameter 6.8 nm, peak diameter 5.4 nm) (**Figure 2B**). Upon reassembly, T4GALA re-forms mostly monodisperse protein cages of the expected diameter (Z-average diameter 76.78 nm, peak diameter 55.31 nm), with a slight increase in aggregation observed by TEM and DLS analysis. (**Figure 2C**).

### Structural characterization of the T4GALA protein nanocage

To further characterize T4GALA, cryogenic electron microscopy (cryo-EM) was carried out on the engineered protein cage. The overall structure of T4GALA shows that it self-assembles into a 7.7 MDa 240mer (T=4) nanocompartment about 42 nm in diameter, nearly identical to native QtEnc (PDB 6NJ8). However, T4GALA exhibits a notable absence of cryo-EM density in the E-loop region between residues Glu58 and Gly83, corresponding to the GALA insertion site **(Figure 3, Figure S5, Figure S6, Table S2)**. Specifically, E-loops at the three-fold symmetry axis formed by three neighboring hexameric capsomers show no density for 21 out of 22 GALA insertion residues – including the glycine linkers. Three additional residues (Glu58, Ser59, and His60) preceding the GALA insertion site lack density as well. At the pseudo-three-fold axis formed by two hexameric and one pentameric capsomer, a similar absence of density is observed around the GALA insertion site (**Figure S7**). While density is visible for all other E-loop residues, model-to-map correlation is relatively low for these E-loop residues across different chains (**Figure S8**), suggesting the engineered E-loop is more structurally dynamic, corroborating the goal of creating a less structurally rigid, triggerable E-loop.

**Figure 3.**
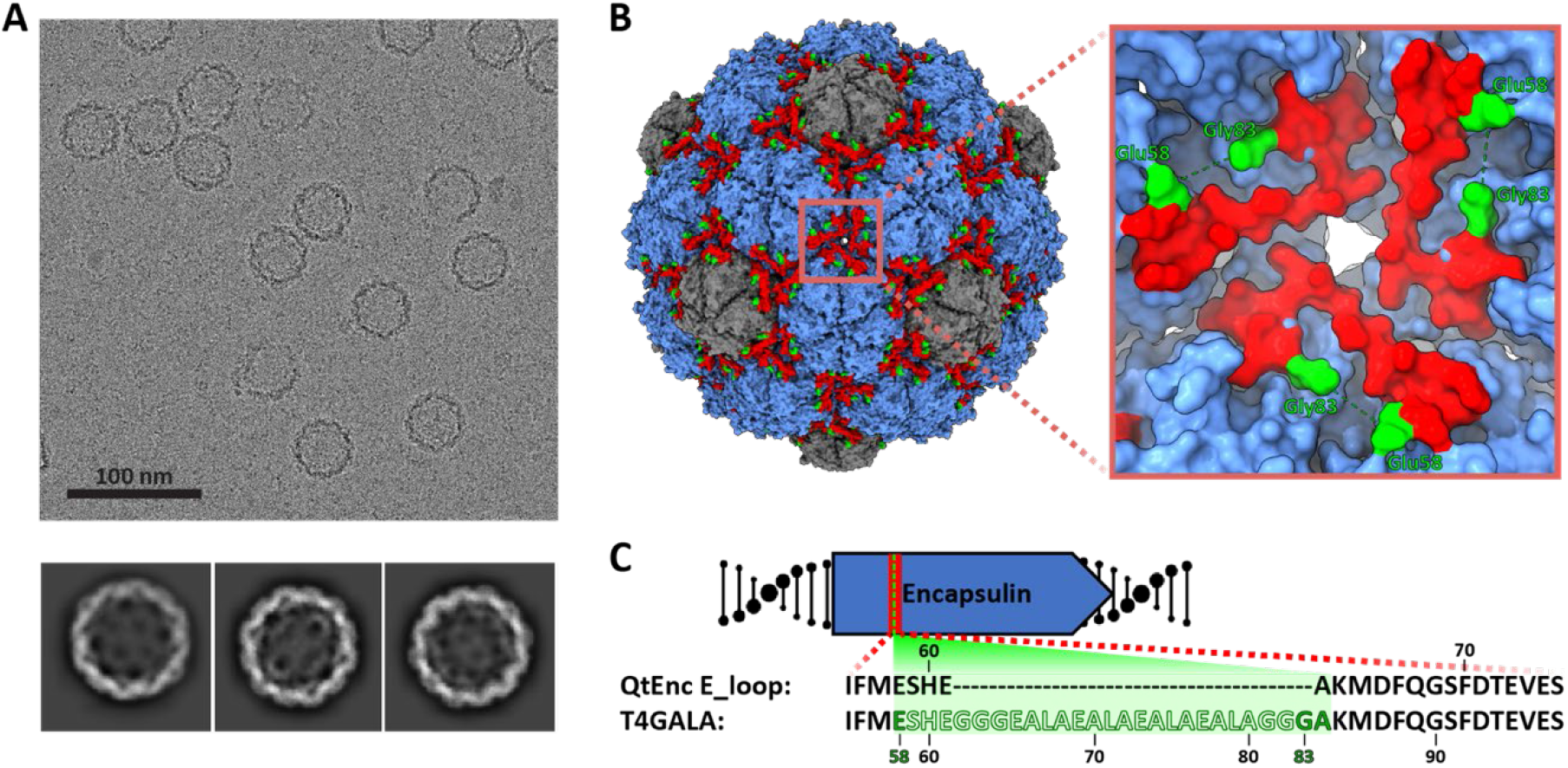
Structural analysis of T4GALA. A) Representative motion-corrected electron cryomicrograph (top) and 2D class averages of T4GALA. B) Cryo-EM density of T4GALA. Hexameric and pentameric capsomers shown in blue and grey, respectively. E-loops are highlighted in red and the last visible residues flanking the GALA insertion site are shown in green (Glu58 and Gly83). Inset (right) highlighting details of the three-fold symmetry axis to emphasize missing E-loop density (Ser59 to Gly82, green dashes). C) Schematic highlighting the observed (solid) and missing (silhouette) residues in the T4GALA E-loop.

### In vivo cargo loading of T4GALA, cargo sequestration, and cargo activity

An N-terminally sumoylated quadruple tandem repeat split fluorescent protein (sFP) was fused at the C-terminus to a QtEnc targeting peptide (T4TP) and cloned immediately upstream of the T4GALA gene for co-expression (**Figure 4A**).^18,27,28^ *In vivo* cargo loading capabilities were then confirmed via Ni-NTA affinity co-purification (**Figure 4B**). Additionally, plate-based sFP complementation fluorescence analysis further confirmed *in vivo* cargo loading while also confirming triggered disassembly capabilities (**Figure 4A, 4B**).^29^ Assembled GFP11_x4_-loaded and disassembled GFP11_x4_-bound T4GALA were individually mixed with separately purified GFP1-10 sFP complement and each separate reaction was allowed to mature overnight for 16 hours. Assembled T4GALA prevented the encapsulated GFP110D7;4 from interacting with GFP1-10 resulting in low relative fluorescence as compared to disassembled T4GALA, which allowed for robust GFP1-10 complementation yielding more than four-fold relative fluorescence. The ability of T4GALA to create a sequestered nanoscale space and robustly encapsulate its cargo until purposefully triggering disassembly will be a useful feature for various biomolecular engineering applications.

**Figure 4.**
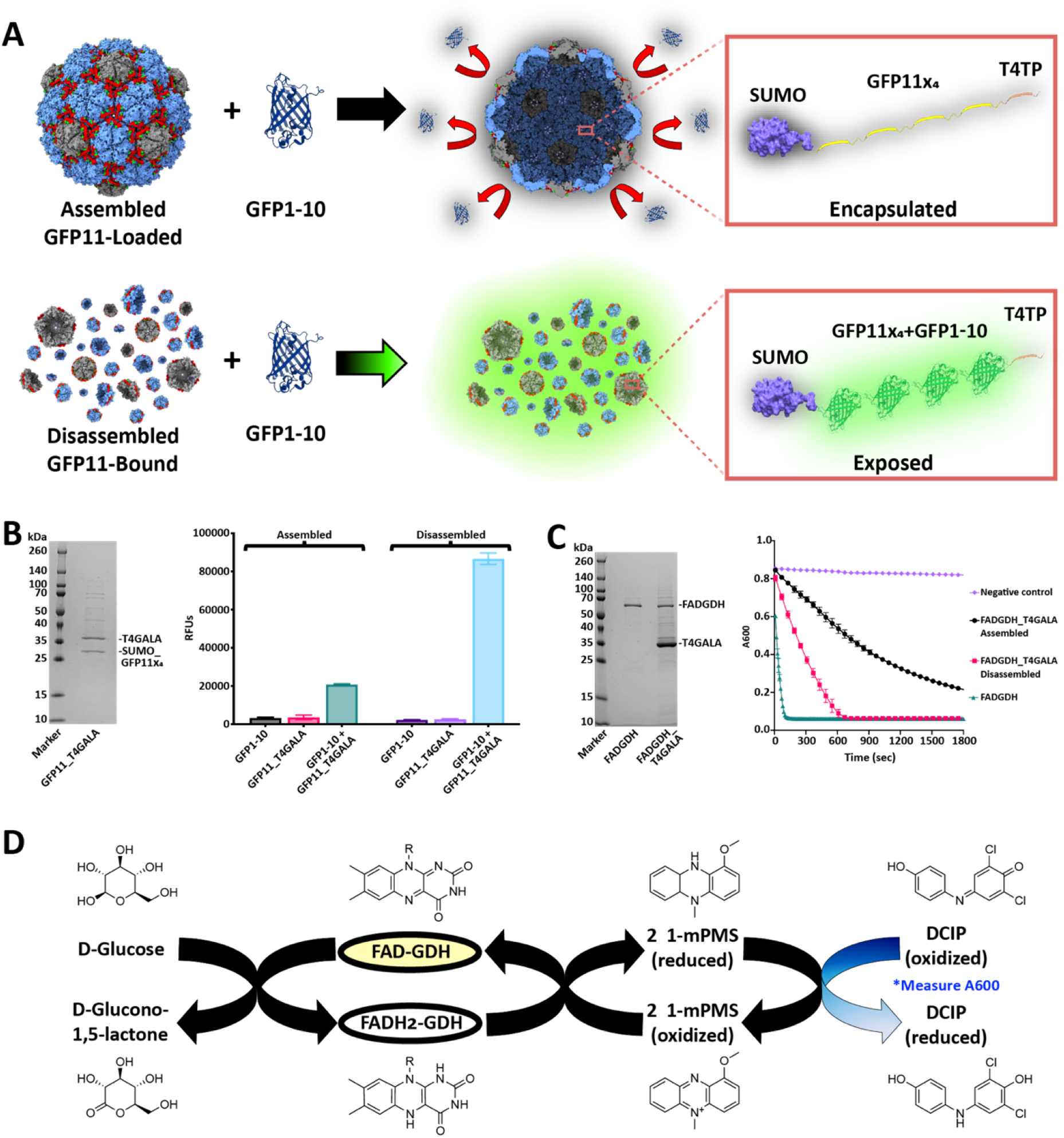
*In vivo* cargo loading of T4GALA and characterization of cargo-loaded systems. A) Schematic of split fluorescent protein experiments. Assembled (top) and disassembled (bottom) GFP11_x4_-loaded/bound T4GALA exposed to the GFP1-10 complement. B) SDS-PAGE analysis of GFP11_x4_-loaded T4GALA (left). Plate-based fluorescence assays (right) showing increased relative fluorescence for disassembled GFP11_x4_-bound T4GALA complementation (light blue; right) compared to roughly four-fold lower fluorescence for an equimolar amount of assembled GFP11_x4_-loaded T4GALA (green, left). C) SDS-PAGE analysis of GDH and GDH-loaded T4GALA (left). Plate-based assays (right) comparing enzymatic activity of unencapsulated FAD-dependent glucose dehydrogenase enzyme (green triangles), *in vivo* T4GALA-encapsulated enzyme in the assembled state (black squares), and *in vivo* T4GALA-encapsulated enzyme in the disassembled state (pink squares) with buffer blank as a negative control (purple diamonds). Data are shown as means while error bars represent standard deviations from three independent experiments. D) Schematic summary of the catalyzed enzymatic reaction and the complementary assay measuring the resultant decrease in absorption at 600 nm as DCIP is reduced. FAD, flavin adenine dinucleotide; GDH, glucose dehydrogenase; 1-mPMS, 1-methoxy-5-methylphenazinium methylsulfate; DCIP, 2,6-dichloroindophenol.

To expand the characterization of *in vivo* loading to enzymes and test potential diffusion barrier effects of encapsulation, a T4 targeting peptide was fused to the C-terminus of a flavin adenine dinucleotide-dependent glucose dehydrogenase enzyme (GDH),^30^ cloned immediately upstream of T4GALA, and co-expressed for *in vivo* encapsulation. Cargo loading capabilities were again confirmed via Ni-NTA affinity co-purification and time-course analyses were conducted via the 2,6-dichloroindophenol (DCIP) assay, which monitors the decrease in absorbance at 600 nm as DCIP is reduced, to determine whether GDH loaded into T4GALA *in vivo* could maintain enzymatic activity (**Figure 4C and 4D**).^30,31^ Comparisons were therefore made between equimolar amounts of free GDH enzyme, encapsulated GDH, and GDH enzyme bound to disassembled T4GALA. While T4GALA-encapsulated GDH exhibited enzymatic activity, the free enzyme displayed substantially faster kinetics. Upon disassembly, the reaction rate increased substantially, but was still observed to be slower than free GDH. It is widely reported throughout the literature that enzymes tethered to a surface often display decreased specific activity,^32^ and it has also been reported that encapsulated enzymes often exhibit decreased specific activity, hypothesized to be the result of rapid *in vivo* encapsulation which may prevent proper folding and cofactor binding.^33^ Additionally, the protein shell likely acts as a diffusion barrier which may decrease the flux of certain substrates and products in and out of the protein nanocage. Therefore, a decrease in encapsulated enzyme activity such as that observed here is not wholly unanticipated. Overall, the *in vivo* encapsulation of an active enzyme, along with its maintained activity after disassembly, highlights the potential modularity and applicability of the T4GALA system.

### *In vitro* cargo loading of T4GALA

To analyze whether the engineered T4GALA protein cage is capable of being disassembled, loaded *in vitro* with exogenous cargo, and then reassembled, a T4 targeting peptide was fused to the C-terminus of mNeonGreen fluorescent protein (mNeonTP). After disassembly of T4GALA, it was mixed with the separately expressed and purified mNeonTP in different molar ratios (6:2 and 6:1 T4GALA:mNeonTP) and then incubated overnight to allow complementation and maturation (**Figure 5A**). Next, T4GALA was reassembled and assessed for *in vitro* cargo loading via native PAGE and fluorescence analysis (**Figure 5b, Figure S9**). Fluorescence of the loaded mNeonTP was observed along with the high molecular weight reassembled T4GALA protein band, suggesting the engineered protein cage is capable of being loaded with exogenous cargo *in vitro*. Importantly, the experiment was conducted in parallel with an alternative mCherry fluorescent protein lacking the T4 targeting peptide as a negative control. The negative control sample failed to exhibit *in vitro* T4GALA encapsulation, indicated by a lack of co-migrating fluorescence during native PAGE analysis. The ability to easily encapsulate proteins inside a defined protein shell under mild conditions *in vitro* once again highlights the potential broad application range of the T4GALA system.

**Figure 5.**
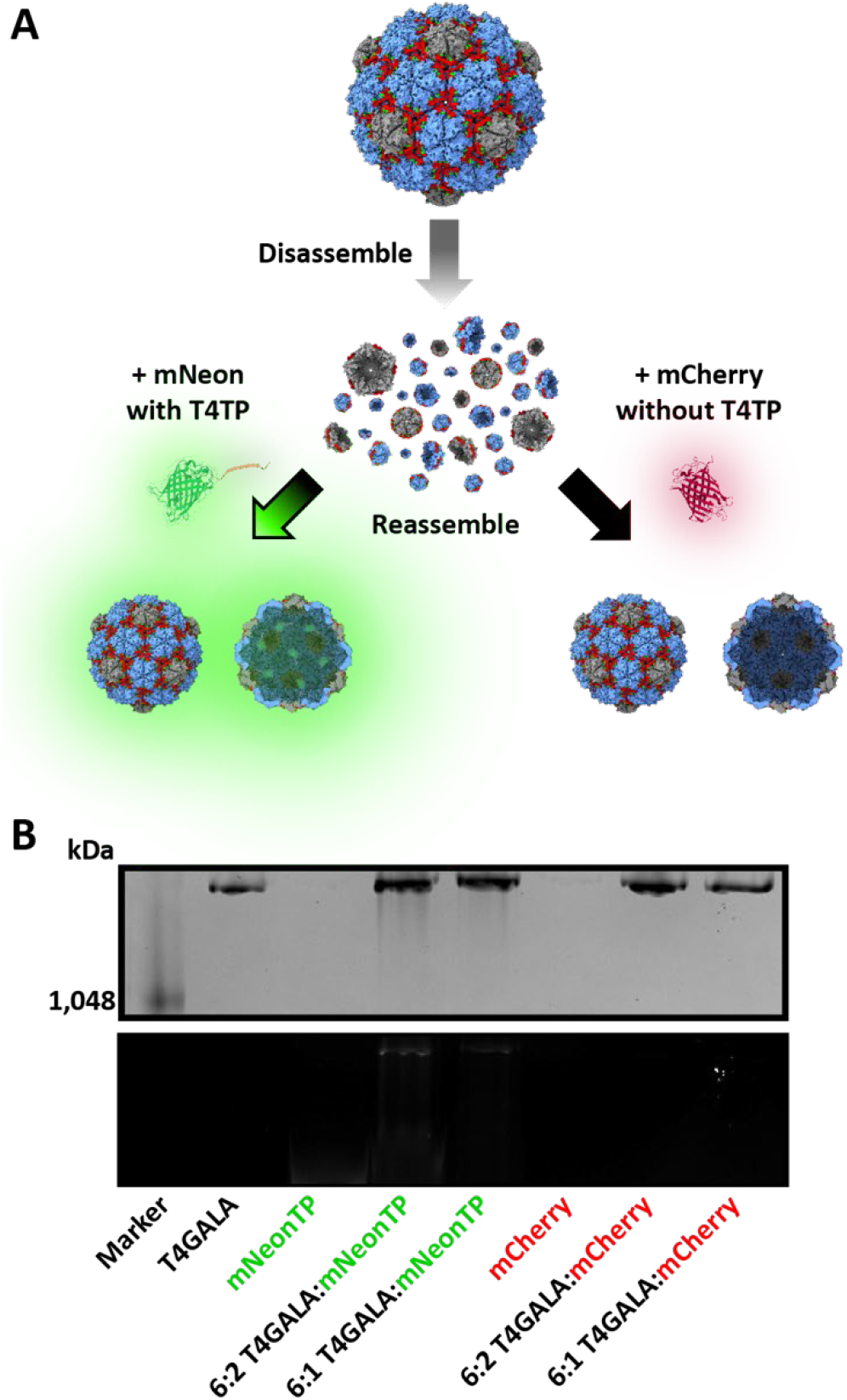
*In vitro* cargo loading of T4GALA. A) Schematic of T4GALA *in vitro* cargo loading including protein cage disassembly, *in vitro* loading of targeting peptide-fused cargo (left) and T4GALA reassembly resulting in detectable fluorescence from newly encapsulated mNeon cargo. Conversely, the same procedure is carried out with mCherry lacking the targeting peptide, which fails to result in cargo loading (right) and results in no detectable fluorescence after reassembly. B) NativePAGE analysis showing high molecular weight bands for assembled T4GALA via Coomassie blue staining (top) and fluorescence analysis of mNeon and mCherry (bottom).

## Conclusion

From bionanoreactors to nanotherapeutic technologies, protein nanocage design presents significant opportunities across numerous research fields. While *de novo* protein cage design has led to several novel biomolecular tools,^34^ increasing numbers of natural protein nanocompartments are being discovered that have been refined by evolution for biological activity and biocompatibility whilst also being amenable to rational engineering approaches.^18,35^ The recent surge in encapsulin nanocompartment discovery and engineering further emphasizes this point.^10,14,36^ Newly discovered protein cages provide an opportunity to create novel semi-synthetic hybrid compartments and bionanotechnological tools. For example, previous research has shown that disassembling and reassembling viral capsids or encapsulins requires extremes of pH^7,37,38^ or salt concentration,^39^ making these manipulations less applicable to biomolecular and biomedical research. In contrast, the T4GALA system described here is functional under milder conditions better suited for conventional experimental procedures and potential biocatalysis or delivery applications. The T4GALA nanocage adds a novel dimension of control to encapsulin nanocages.

Via simple buffer exchanges within physiological pH and ionic strength ranges, the T4GALA system showcases the ability to undergo on-demand disassembly and reassembly. Structural analyses via cryo-EM confirm our overall design strategy by highlighting a lack of density for the rationally engineered disassembly trigger and an altogether more dynamic E-loop. The engineered protein cage also retains the ability of *in vivo* cargo loading via co-expression with targeting peptide-fused proteins of choice. Additionally, facile *in vitro* cargo loading under mild conditions represents a novel capability for encapsulin nanocages.

Potential applications of the T4GALA system include control over the unloading of relatively large encapsulated nanoreactor products, sequentially timed exposure of protected cargos to external molecules, *in vitro* encapsulation of enzymes that cannot be co-expressed with T4GALA, or even stoichiometric shuffling of nanocage components. In sum, the T4GALA system developed here represents a versatile addition to the growing encapsulin-based biomolecular engineering toolbox.

## Data Availability

The determined structure has been deposited and the model was assigned the accession code PDB ID 7MH2. The final cryo-EM map was submitted to EMDB with the identifier 23834. All other data that support the findings of this study are available from the corresponding author upon request.

## Acknowledgements

We gratefully acknowledge funding from the NIH (1R35GM133325). Research reported in this publication was supported by the University of Michigan Cryo-EM Facility (U-M Cryo-EM). U-M Cryo-EM is grateful for support from the U-M Life Sciences Institute and the U-M Biosciences Initiative.

Molecular graphics and analyses performed with UCSF ChimeraX, developed by the Resource for Biocomputing, Visualization, and Informatics at the University of California, San Francisco, with support from National Institutes of Health R01-GM129325 and the Office of Cyber Infrastructure and Computational Biology, National Institute of Allergy and Infectious Diseases.

## Author Contributions

A.S.C., J.A.J., and T.W.G. designed the project. A.S.C. and T.W.G. designed the engineered protein cage. A.S.C. and J.A.J. conducted the laboratory experiments and transmission electron microscopy, while M.P.A. obtained and analyzed cryo-electron microscopy data. T.W.G. oversaw the project in its entirety.

## Competing Interests

The authors declare no competing interests.

## Supporting Information

Supporting Information containing methods and additional data and analyses is available and contains Figures S1-S9 and Tables S1-S2.

## Notes

### Competing Interest Statement

The authors have declared no competing interest.

