## Supplementary Information for "Triggered reversible disassembly of an engineered protein nanocage"

**Abstract**

Protein nanocages play crucial roles in sub-cellular compartmentalization and spatial control in all domains of life and have been used as biomolecular tools for applications in biocatalysis, drug delivery, and bionanotechnology. The ability to control their assembly state under physiological conditions would further expand their practical utility. To gain such control, we introduced a peptide capable of triggering conformational change at a key structural position in the largest known encapsulin nanocompartment. We report the structure of the resulting engineered nanocage and demonstrate its ability to on-demand disassemble and reassemble under physiological conditions. We demonstrate its capacity for *in vivo* encapsulation of proteins of choice while also demonstrating *in vitro* cargo loading capabilities. Our results represent a functionally robust addition to the nanocage toolbox and a novel approach for controlling protein nanocage disassembly and reassembly under mild conditions.

### 1. Materials

#### 1.1 Chemical and biological materials

All chemicals were used as supplied by vendors without further purification. Imidazole, Invitrogen Novex WedgeWell 14% Tris-glycine Mini Protein Gels, Isopropy-ß-D-thiogalactopyranoside (IPTG), lysozyme, NativePAGE™ 4 to 16% bis-tris mini protein gels, NativeMark Unstained Protein Standard, Spectra™ Multicolor Broad Range Protein Ladder, Thermo Scientific Pierce 660nm Protein Assay Reagent, Tris base, Tris HCl, all restriction enzymes, and all cell culture media and reagents were purchased from Fisher Scientific, Inc. (USA). Gibson Assembly Master Mix was purchased from NEB (USA). Amicon Ultra-0.5 mL centrifugal units and Benzonase® nuclease were purchased from MilliporeSigma (USA). BL21 (DE3) Electrocompetent Cells used for *E. coli* expression were also purchased from MilliporeSigma (USA). Bis-tris propane from Research Products International (USA) was used for the assembly buffer. Ni-NTA agarose from Gold Biotechnology, Inc. (USA) was used for His-tagged protein purification.

#### 1.2 Instrumentation

Lysate sonication was conducted with a Model 120 Sonic Dismembrator from Fisher Scientific, Inc. (USA). Protein quantification was carried out using a Nanodrop Spectrophotometer from ThermoFisher Scientific, Inc. (USA) and a Synergy H1 Microplate Reader BioTek Instruments (USA) as applicable. All size-exclusion chromatography (SEC) was carried out on an AKTA Pure fast liquid protein chromatography system with a Superose 6 10/300 GL (Cytiva, USA). Polyacrylamide gel electrophoresis (PAGE) and NativePAGE were performed in an XCell SureLock from Invitrogen/ThermoFisher Scientific (USA). Gel images were captured using a ChemiDoc Imaging System from Bio-Rad Laboratories, Inc. (USA). DLS was carried out on an Uncle from Unchained Labs (USA). TEM was carried out on a Morgagni 100 keV Transmission Electron Microscope (FEI, USA). Plate-based absorbance and fluorescence assays were conducted on the Synergy H1 Microplate Reader BioTek Instruments (USA). Glow discharging was conducted with a PELCO easiGlow™ system by Ted Pella, Inc (USA). A Glacios Cryo Transmission Electron Microscope by ThermoScientific, Inc. (USA) equipped with a K2 Summit direct electron detector by Gatan, Inc. (USA) was used for cryo-EM. Smaller materials are listed along with corresponding methods below.

#### 1.3 Software

The following software was used throughout this work: Adobe Illustrator 2021 v25.0.0 (figures), CryoSPARC v2.15.00[1] (cryo electron microscopy), Fiji/ImageJ v2.1.0/1.53c[2] (densitometric data analysis and TEM images), GraphPad Prism for Mac OS v9.0.2 (fluorescence assay graphs), Bio-Rad Image Lab Touch Software (gel imaging), Microsoft Excel for Mac v16.46 (DLS graphs), Phenix v1.18.2-3874[3] (model building), UCSF Chimera v1.14[4] and ChimeraX v1.1[5] (protein structure), and UNICORN 7 (FPLC system control and chromatography).

### 2. Experimental Methods

#### 2.1 Sequence alignments

The QtEnc and T4GALA alignment was generated with the ESPript 3 server (http://espript.ibcp.fr/) using a protein sequence alignment produced with Clustal Omega, with secondary structure information extracted from the original QtEnc structure (PDB 6NJ8; **Figure S1**).[6,7]

#### 2.2 Protein production

QtEnc was expressed and purified as previously described.[8] For all other proteins, plasmids were either constructed in lab with target gBlock genes codon-optimized and synthesized by IDT (USA) and inserted into the pETDuet-1 vector via Gibson assembly around the NdeI and PacI restriction sites, or synthesized and inserted into the pET-28a(+) vector by Twist Bioscience (USA) (**Table S1)**. Plasmids were transformed into BL21 (DE3) *E. coli* cells via electroporation per protocol and 25% glycerol bacterial stocks were made and stored at 80°C until further use. Starter cultures were grown in 5 mL LB with the appropriate selection antibiotic (either 100 mg/mL ampicillin or 50 mg/mL kanamycin) at 37°C overnight. For all constructs containing the T4GALA protein nanocage (either empty or cargo-loaded), 500 mL of LB with the appropriate selection antibiotic were inoculated with overnight starter cultures and grown at 37°C to an OD600 of 0.4-0.5, then induced with 0.1 mM IPTG and grown further at 30°C overnight for ~18 hours. For un-encapsulated FADGDH, GFP1-10, mCherry, and mNeonTP proteins, 500 mL of LB with the appropriate selection antibiotic was inoculated with the respective overnight starter culture and grown at 37°C to OD600 ~0.6, then induced with 0.5 mM IPTG and grown for ~18 additional hours overnight at 18°C. Cells were then harvested via centrifugation at 10,000 rcf x 15 minutes at 4°C and pellets were frozen and stored at -80°C until further use.

#### 2.3 Protein purification

Other than QtEnc, all proteins were initially purified via Ni-NTA batch purification. Briefly, frozen pellets were thawed on ice and resuspended in ~5 mL/g (wet cell mass) of cold Resuspension Buffer (50 mM Tris pH 8.0, 150 mM NaCl, 1 mM TCEP, 10 mM imidazole, and 5% glycerol). Lysis components were added (0.5 mg/mL lysozyme, one SIGMA*FAST* EDTA-free protease inhibitor cocktail tablet per 100 mL, 0.5 mM $MgCl_2$, and 25 units/mL Benzonase® nuclease) and samples were lysed on ice for 10 minutes. Samples were then sonicated at 50% amplitude for 5 minutes total (eight seconds on, 16 seconds off) until no longer viscous. After sonication, samples were centrifuged at 8,000 rcf x 15 minutes at 4°C. Lysate was then bound to Ni-NTA resin pre-equilibrated with Resuspension Buffer via rocking at 4°C for 45 minutes. Supernatant was discarded and the bound sample was washed once with normal Resuspension Buffer and a second time with Resuspension Buffer with 20 mM imidazole. Protein was then eluted three times with Elution Buffer (50 mM Tris pH 8.0, 150 mM NaCl, 1 mM TCEP, 350 mM imidazole, 5% glycerol) and either stored at 4°C for future use or, as in the case of T4GALA, immediately purified further via SEC. For SEC purification, post-Ni-NTA eluted T4GALA target protein was immediately centrifuged at 10,000 rcf x 10 minutes, then loaded on an AKTA Pure and purified via Superose 6 10/300 GL pre-equilibrated with Elution Buffer. All proteins were stored at 4°C until use.

#### 2.4 Preliminary assembly and disassembly buffer analyses

T4GALA was purified via batch Ni-NTA and the third elution was used for preliminary assembly/disassembly trials. T4GALA at 0.4 mg/mL was buffer exchanged into various buffers containing 150 mM NaCl ranging from pH 5.5 to pH 8.0 in 0.5 pH unit increments (**Figure S2**) via five successive exchanges in 100 kDa MWCO Amicon Ultra-0.5 mL centrifugal units. Larger molecular weight cutoff filters were used to assure collection of assembled encapsulin while allowing disassembled T4GALA to pass through the filter. This was done out of concern that re-collected disassembled T4GALA could potentially reassemble from extended exposure to the NativePAGE running buffer due to relatively higher pH (pH 8.3) and the designed pH-sensitivity of the disassembly trigger. Preliminary SEC trials were also conducted to analyze T4GALA assembly state during SEC purification in different buffers to assess assembly dependence on imidazole ionic strength (Resuspension Buffer vs Elution Buffer; **Figure S3**).

#### 2.5 Native polyacrylamide gel electrophoresis

All Native PAGE analyses were conducted in an Invitrogen XCell SureLock using NativePAGE™ 4 to 16% bis-tris mini protein gels and NativeMark Unstained Protein Standard from Fisher Scientific (USA) with 1x running buffer made from 10x Tris/Glycine Buffer from Bio-Rad Laboratories, Inc. (USA). 10 µg of protein was loaded per well, with effort to maintain equivalent amounts across all lanes for comparative analyses. Native PAGE gels were run overnight at 65V for 16.5 hours at 4°C. The following day, gels were stained with Instant*Blue*™ from Expedeon Ltd. (UK) and imaged and analyzed.

#### 2.6 Disassembly and reassembly of the engineered protein nanocage

T4GALA was rapidly purified in Elution Buffer in the assembled state (**Figure 2A**). The sample was then disassembled by buffer exchanging into Disassembly Buffer (20 mM Tris pH 7.1, 150 mM NaCl) via five successive exchanges in 30 kDa MWCO Amicon Ultra-0.5 mL centrifugal units and incubating at 25°C for at least six hours. Sample was then either stored at 4°C until further use (**Figure 2B**) or reassembled. For reassembly, the sample was buffer exchanged as before, but into Reassembly Buffer (20 mM bis-tris propane pH 7.5, 150 mM NaCl, 25 mM imidazole) with substantial pipette mixing and incubated at 25°C overnight for ~16 hours. Sample was then stored at 4°C until further use (**Figure 2C**).

#### 2.7 Dynamic light scattering analysis

All sizing and polydispersity measurements were carried out on an Uncle by Unchained Labs (USA) at 30 °C in triplicate. Purified T4GALA samples were adjusted to 0.4 mg/mL of monomer in the appropriate corresponding buffers and centrifuged at 10,000 rcf x 10 minutes, then immediately analyzed via DLS in either the assembled state (**Figure 2A**), the disassembled state (**Figure 2B**), or in the reassembled state (**Figure 2C**).

#### 2.8 Transmission electron microscopy

Negative stain transmission electron microscopy (TEM) was carried out on the various T4GALA samples (0.4 mg/mL), FADGDH_T4GALA (0.1 mg/mL), and GFP11_T4GALA (0.4 mg/mL) with 200-mesh gold grids coated with extra thick (25-50 nm) formvar-carbon film (EMS, USA) made hydrophilic by glow discharging at 5 mA for 60 s. Briefly, 3.5 µL of sample was added to the grid and incubated for one minute, wicked with filter paper, and washed once with distilled water and once with 0.75% (w/v) uranyl formate before staining with 8.5 µL of uranyl formate for one minute. TEM images were captured using a Morgagni transmission electron microscope at 100 keV at the University of Michigan Life Sciences Institute. For all TEM experiments, samples were at 0.35 mg/mL of T4GALA monomer in appropriate buffer.

#### 2.9 Cryo-EM sample preparation

Purified T4GALA protein cage sample was buffer exchanged into 20 mM bis-tris propane pH 7.5, 150 mM NaCl and concentrated to 0.76 mg/ml. 3.5 µL of sample was applied to glow discharged grids (Quantifoil R1.2/1.3 grids, glow discharged 1 minute, 5 mA) which were frozen by plunging into liquid ethane using an FEI Vitrobot Mark IV (100% humidity, temperature: 22 °C, blot force: 20, blot time: 4 seconds, wait time: 0 seconds). Grids were clipped and stored in liquid nitrogen until data collection.

#### 2.10 Cryo-EM data acquisition

Cryo-EM movies were collected using a ThermoFisher Scientific Glacios cryo-electron microscope operating at 200 kV equipped with a Gatan K2 Summit direct electron detector located at the University of Michigan Life Sciences Institute. Movies were collected at 45,000 magnification with a pixel size of 0.98 Å using Leginon software package with exposure time of 8 seconds, frame time of 200 ms, and total dose of 62 $e^-/Å^2$.[9] 1,713 movies were collected in a single session on August 20, 2020.

#### 2.11 Cryo-EM data processing

All data processing was performed using CryoSPARC V 2.15.00.[1] Movies were imported, and motion corrected using Patch motion correction. Patch CTF estimation was used for CTF correction. Motion and CTF corrected movies were then curated using Curate Exposures to select only movies with CTF fit resolutions higher than 6 Å, resulting in 1490 exposures to be used downstream. 314 particles were picked using Manual picker and assigned into initial 2D class averages using 2D class with 10 classes. Four good classes were used as templates for Template picker, and selected particles were curated using Inspect Picks to manually identify an optimal NCC and local power score. 11,194 particles were initially extracted from 1,259 exposures with a box size of 630 pixels. Extracted particles were then used as inputs for two successive rounds of 2D classification using 100 and 50 classes with default settings, resulting in 8,300 good particles selected from four classes. Particles were then further filtered by 3D Heterogenous Refinement the *Quasibacillus thermotolerans* T4 encapsulin structure (PDB 6NJ8) as the initial model with a particle size of 256 voxels and I symmetry applied.[8] 6,707 particles from the largest class were down sampled to a box size of 440 pixels and refined via Homogenous Refinement (I symmetry) followed by Local and Global CTF refinements with default settings. This was repeated three times and followed by a final round of 3D refinement giving a final map with a corrected GSFCS resolution of 3.57 Å.

#### 2.12 Model building

The final cryoSPARC map was opened in UCSF Chimera v 1.14.[4] An initial model (PDB 6NJ8) containing four protomers was placed into the density by manually aligning to the map followed by a rigid body fit using the fit to volume command. Coordinates were opened in Coot v 0.9-pre and manually refined against the map initially with a rigid body refinement, followed by chain refinement, and iterative manual real space refinements.[10,11] This was done for each of the 4 chains in the asymmetric unit (ASU) of the model. The model was then refined in Phenix v 1.18.2-3874 using phenix.real_space_refine with default parameters for the ASU.[3,12,13] Non-crystallographic symmetry (NCS) was identified in the cryo-EM map using phenix.map_symmetry, and NCS operators were applied using phenix.apply_ncs to make a complete model containing 60 copies of the ASU in 240 different chains. This model was then refined using phenix.real_space_refine with NCS restraints, global minimization, and ADP refinements. The model was validated using the Comprehensive Validation tool via phenix.validation_cryoem. The model was inspected and manually refined until satisfactory geometry and density fit statistics were reached. Prior to depositing the structures to RCSB databank, the models were again validated using phenix.validation_cryoem (**Figure 3, Figure SS4, Figure S5, Figure S6**). The model was deposited to the PDB under PDB ID 7MH2 and the EMDB under EMD-23834.

#### 2.13 Fluorescence-based assays

T4GALA with encapsulated sumoylated four-fold tandem repeat of GFP11 split fluorescent protein (sFP) fused to a T4 targeting peptide (GFP11_T4GALA) was purified separately from His-tagged GFP1-10 sFP. Then, GFP11_T4GALA was split into two samples and one was buffer exchanged into Disassembly Buffer while the other was buffer exchanged into Reassembly Buffer. Next, each GFP11_T4GALA sample was separately mixed at 1 µM with 5 µM GFP1-10 in the appropriate buffers and allowed to complement and mature at 25°C overnight for ~16 hours in duplicate. Endpoint GFP1-10/11 fluorescence (485 nm/510 nm) was then then measured in a BioTek Synergy H1 microplate reader at a final volume of 100 µL in Corning® 96-well flat clear bottom black polystyrene microplates (**Figure 4**). Additionally, T4GALA was purified, concentrated (12 µM ), and disassembled, then mixed with two different concentrations (2 µM  and 4 µM) of separately Ni-NTA purified mNeon fluorescent protein fused to a T4 targeting peptide (mNeonTP). Samples were then reassembled and analyzed via NativePAGE analysis for fluorescence in the presence of the higher assembled protein band after blue staining. The experiment was conducted in parallel with mCherry that lacked a targeting peptide, and the two samples were compared.

#### 2.14 Glucose dehydrogenase activity assays

Flavin adenine dinucleotide-dependent glucose dehydrogenase (FADGDH) from *Rasamonia emersonii* (previously *Taloromyces emersonii*) was purified in both T4GALA-encapsulated and unencapsulated, free states. Equimolar free FADGDH, assembled T4GALA-encapsulated FADGDH, and disassembled T4GALA-bound FADGDH were analyzed to analyze enzyme activity via the 2,6-dichloroindophenol (DCIP) assay method. Briefly, the reaction buffer (140 µL) contained either 50 mM bis-tris propane (assembled T4GALA-encapsulated FADGDH assay) or 50 mM tris (disassembled T4GALA-bound FADGDH assay) at pH 7.5, with 0.14 mM 1-methoxy-5-methylphenazinium methylsulfate (1-mPMS), 0.07 mM DCIP sodium salt hydrate, and 300 mM glucose. Reactions were started with the addition of 10 µL of the various enzyme samples (or buffer blanks) to bring the reaction to a final concentration of 100 nM FADGDH enzyme. The reduction of DCIP was measured at 600 nm in a BioTek Synergy H1 microplate reader at a final volume of 150 µL in Corning® 96-well flat clear bottom black polystyrene microplates in 10 second intervals for a total of 30 minutes to evaluate enzyme activity and the rate of reaction. Free, unencapsulated FADGDH assays were conducted in duplicate with all other assays carried out in triplicate.

### 3. Supporting Results and Schematic Diagrams

#### 3.1 Design of the protein nanocage and the disassembly trigger

The protein nanocage was designed by inserting the 16 residue EALA quadruple repeat GALA peptide between flanking triple glycine linkers. The peptide was inserted within the E-loop between Glu61 and Ala62.

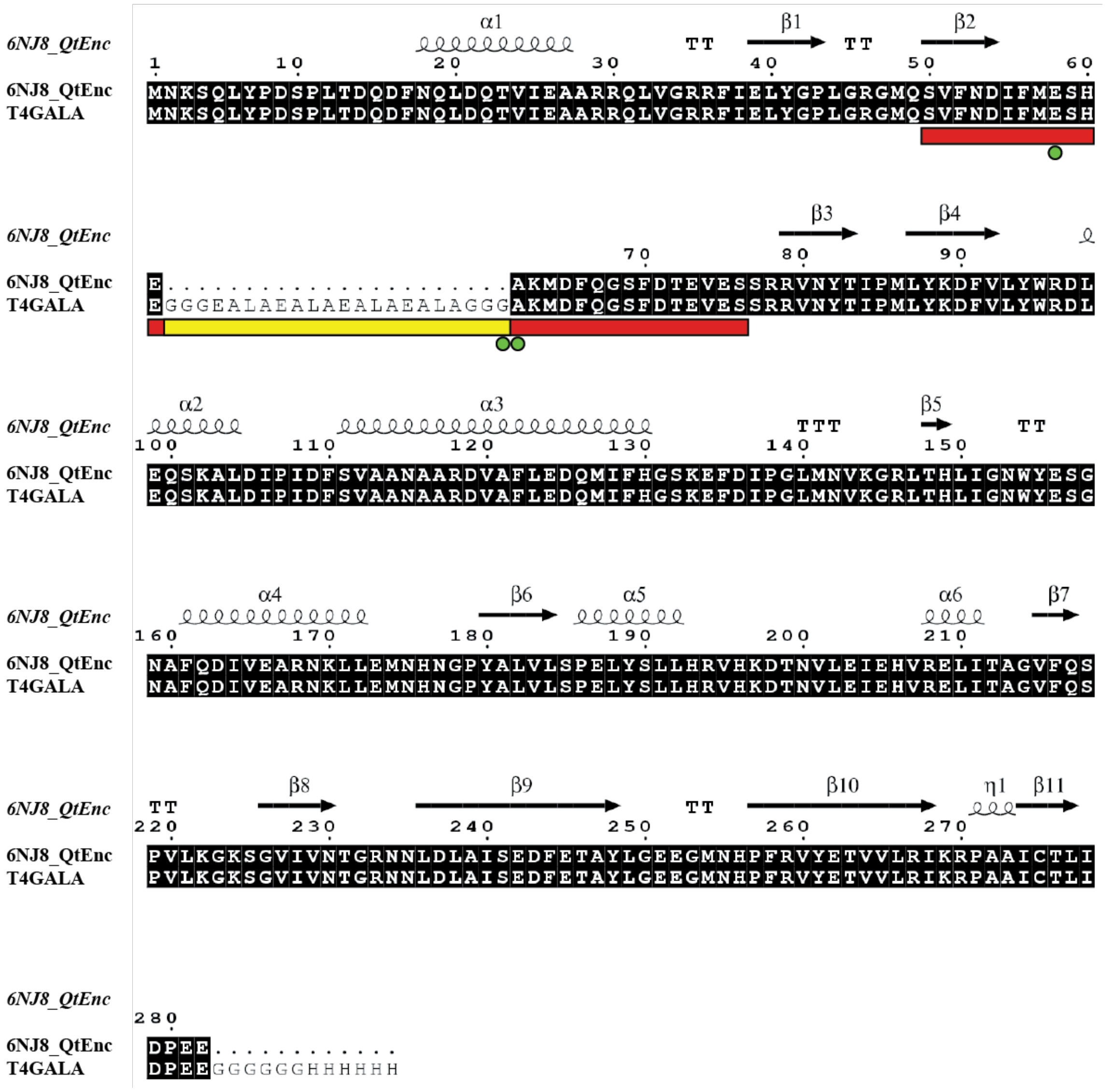

**Figure S1.** Sequence alignment of the encapsulin from *Quasibacillus thermotolerans* (QtEnc; PDB 6NJ8) and the T4GALA engineered protein nanocage. T4GALA was created from the T=4 QtEnc by inserting a 16 residue GALA repeat with triple glycine flanking repeats between residues E61 and A62 located within the QtEnc E-loop. Identical aligned residues are highlighted in black, the E-loop is indicated with a red bar, the GALA insert is indicated with a yellow bar, and first flanking residues around the three-fold and pseudo-three-fold axes with observable density on cryo-EM analysis indicated with green dots (Figure 3 and Figure S5).

#### 3.2 Design of experimental constructs

In order to fully analyze the assembly, triggered disassembly, and reassembly capabilities of T4GALA, as well as the *in vivo* and *in vitro* cargo loading capabilities of T4GALA, a number of experimental constructs were created as follows.

**Table S1.** Protein sequences of constructs used.

| Construct | Protein sequence[a][b] |
|---|---|
| T4GALA | MNKSQLYPDSPLTDQDFNQLDQTVIEAARRQLVGRRFIELYGPLGRGMQSVFNDIFME<br>SHEGGGEALAEALAEALAEALAGGGAKMDFQGSFDTEVESSRRVNYTIPMLYKDFVLY<br>WRDLEQSKALDIPIDFSVAANAARDVAFLEDQMIFHGSKEFDIPGLMNVKGRLTHLIGN<br>WYESGNAFQDIVEARNKLLEMNHNGPYALVLSPELYSLLHRVHKDTNVLEIEHVRELITA<br>GVFQSPVLKGKSGVIVNTGRNNLDLAISEDFETAYLGEEGMNHPFRVYETVVLRIKRPA<br>AICTLIDPEE* |
| GFP11_T4GALA | MSDSEVNQEAKPEVKPEVKPETHINLKVSDGSSEIFFKIKKTTPLRRLMEAFAKRQGKE<br>MDSLRFLYDGIRIQADQTPEDLDMEDNDIIEAHREQIGGGGSGGSGGSRDHMVLHEY<br>VNAAGITGGSGGSGGSGGRDHMVLHEYVNAAGITGGSGGSGGSGGRDHMVLHEYV<br>NAAGITGGSGGSGGSGGRDHMVLHEYVNAAGITGGSGGSGGSGGHKKKGFTVGSLI<br>Q*…MNKSQLYPDSPLTDQDFNQLDQTVIEAARRQLVGRRFIELYGPLGRGMQSVFND<br>IFMESHEGGGEALAEALAEALAEALAGGGAKMDFQGSFDTEVESSRRVNYTIPMLYKD<br>FVLYWRDLEQSKALDIPIDFSVAANAARDVAFLEDQMIFHGSKEFDIPGLMNVKGRLTH<br>LIGNWYESGNAFQDIVEARNKLLEMNHNGPYALVLSPELYSLLHRVHKDTNVLEIEHVR<br>ELITAGVFQSPVLKGKSGVIVNTGRNNLDLAISEDFETAYLGEEGMNHPFRVYETVVLRI<br>KRPAAICTLIDPEE* |
| GFP1-10 | MGSSHHHHHHGGSENLYFQGMSKGEELFTGVVPILVELDGDVNGHKFSVRGEGEGD<br>ATIGKLTLKFICTTGKLPVPWPTLVTTLTYGVQCFSRYPDHMKRHDFFKSAMPEGYVQE<br>RTISFKDDGKYKTRAVVKFEGDTLVNRIELKGTDFKEDGNILGHKLEYNFNSHNVYITAD<br>KQKNGIKANFTVRHNVEDGSVQLADHYQQNTPIGDGPVLLPDNHYLSTQTVLSKDPNE<br>K* |
| FADGDH | MGSSSYDYIVVGGGTSGLVVANRLSENPNVSVLVIEAGDSVYNNANVTNVNGYGLAFG<br>TPIDWQYQSTNQTYAGNTRQTLRAGKALGGTSTINGMAYTRAQDVQIDAWAAIGND<br>GWDWSSLWPYYLKSEAFTAPNQTQRAAGASYNPAYHGVTGPLHVGFIEMQPNNLSSI<br>LNQTYQALGVPWTEDVNGGKMRGYNFFPSTVDDAADVREDAARAYYYPFESRPNLR<br>VMLNTLANRIVWKNETSGGNVTADGVEVTPLNGTVCRIQANNEVILSAGSLRSPGILEL<br>SGVGNPSILNKYNIPVKVNLPTVGENMQDQMYNDASAEGYSAIAGTKSVAYPSVTDLF |

GNRTSAVAASVQNQLAQYAEAAANQSQGTMKASDLQRLFQIQYDLIFKQEVPIAEIITY
PTGNTLAAGYWGLLPFARGSVHIASADPTVQPVINPNYFMFDWDVQQQIGTAKFIRN
LYKTAPLSSLVKNETEPGSAVPEGASDSVWEAWLKETYRSNFHPVGTAAIMPRSIGGVV
DERLRVYGTANVRVVDASVLPFQICGHLTSTLYAVAERASDFLKEDAARLGNGGSGGSE
NLYFQGGSGGSGGSHHHHHH*

FADGDH_T4GALA

MGSSSYDYIVVGGGTSGLVVANRLSENPNVSVLVIEAGDSVYNNANVTNVNGYGLAFG
TPIDWQYQSTNQTYAGNTRQTLRAGKALGGTSTINGMAYTRAQDVQIDAWAAIGND
GWDWSSLWPYYLKSEAFTAPNQTQRAAGASYNPAYHGVTGPLHVGFIEMQPNNLSSI
LNQTYQALGVPWTEDVNGGKMRGYNFFPSTVDDAADVREDAARAYYYPFESRPNLR
VMLNTLANRIVWKNETSGGNVTADGVEVTPLNGTVCRIQANNEVILSAGSLRSPGILEL
SGVGNPSILNKYNIPVKVNLPTVGENMQDQMYNDASAEGYSAIAGTKSVAYPSVTDLF
GNRTSAVAASVQNQLAQYAEAAANQSQGTMKASDLQRLFQIQYDLIFKQEVPIAEIITY
PTGNTLAAGYWGLLPFARGSVHIASADPTVQPVINPNYFMFDWDVQQQIGTAKFIRN
LYKTAPLSSLVKNETEPGSAVPEGASDSVWEAWLKETYRSNFHPVGTAAIMPRSIGGVV
DERLRVYGTANVRVVDASVLPFQICGHLTSTLYAVAERASDFLKEDAARLGNGGSGGSG
GSGGHKKKGFTVGSLIQ*…MNKSQLYPDSPLTDQDFNQLDQTVIEAARRQLVGRRFIE
LYGPLGRGMQSVFNDIFMESHEGGGEALAEALAEALAEALAGGGAKMDFQGSFDTEV
ESSRRVNYTIPMLYKDFVLYWRDLEQSKALDIPIDFSVAANAARDVAFLEDQMIFHGSK
EFDIPGLMNVKGRLTHLIGNWYESGNAFQDIVEARNKLLEMNHNGPYALVLSPELYSLL
HRVHKDTNVLEIEHVRELITAGVFQSPVLKGKSGVIVNTGRNNLDLAISEDFETAYLGEE
GMNHPFRVYETVVLRIKRPAAICTLIDPEE*

mNeonTP

MGSSHHHHHHGGSENLYFQGMVSKGEEDNMASLPATHELHIFGSINGVDFDMVGQ
GTGNPNDGYEELNLKSTKGDLQFSPWILVPHIGYGFHQYLPYPDGMSPFQAAMVDGS
GYQVHRTMQFEDGASLTVNYRYTYEGSHIKGEAQVKGTGFPADGPVMTNSLTAADW
CRSKKTYPNDKTIISTFKWSYTTGNGKRYRSTARTTYTFAKPMAANYLKNQPMYVFRKT
ELKHSKTELNFKEWQKAFTDVMGMDELYKKKGFTVGSLIQ*

mCherry

MHHHHHHGGSVSKGEEDNMAIIKEFMRFKVHMEGSVNGHEFEIEGEGEGRPYEGTQ
TAKLKVTKGGPLPFAWDILSPQFMYGSKAYVKHPADIPDYLKLSFPEGFKWERVMNFE
DGGVVTVTQDSSLQDGEFIYKVKLRGTNFPSDGPVMQKKTMGWEASSERMYPEDGA
LKGEIKQRLKLKDGGHYDAEVKTTYKAKKPVQLPGAYNVNIKLDITSHNEDYTIVEQYER
AEGRHSTGGMDELYK*

---

[a] Intergenic sequences for encapsulated constructs represented by ellipses. [b] Stops indicated by asterisks.

### 3.3 T4GALA assembly and disassembly analyses via Native PAGE

The assembly state of T4GALA was initially assessed by fully exchanging into various buffers. Assembly conditions were determined to be dependent upon pH, ionic strength, and buffer identity (**Figure S2**). In summary, buffer conditions resulting in assembly included higher pH, higher imidazole, ionic strength, and bis-tris propane buffer. Conversely, disassembly was driven by lower pH, absence or lower concentrations of imidazole, and tris buffer.

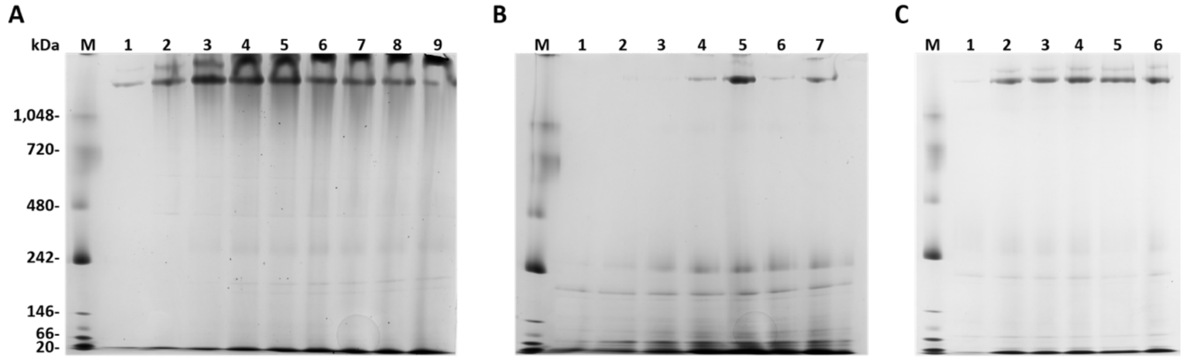

**Figure S2.** Native PAGE analyses of assembly and disassembly of T4GALA. **A**) Native PAGE analysis of QtEnc across a wide pH range for initial comparison shows marked stability and suggests QtEnc stays assembled across a wide pH range. M, marker; 1, 20 mM citrate pH 4.5, 150 mM NaCl; 2, 20 mM citrate pH 5.0, 150 mM NaCl; 3, 20 mM MES pH 5.5, 150 mM NaCl; 4, 20 mM MES pH 6.0, 150 mM NaCl; 5, 20 mM MES pH 6.5, 150 mM NaCl; 6, 20 mM bis-tris propane pH 7.0, 150 mM NaCl; 7, 20 mM tris pH 7.5, 150 mM NaCl; 8, 20 mM Tris pH 8.0, 150 mM NaCl; 9, 20 mM Tris pH 8.5, 150 mM NaCl. **B**) Native PAGE analysis of T4GALA across a wide pH range shows assembly state is substantially dependent upon pH and actual buffer chemical. M, marker; 1, 20 mM citrate pH 5.0, 150 mM NaCl; 2, ; 2, 20 mM MES pH 5.5, 150 mM NaCl; 3, 20 mM MES pH 6.0, 150 mM NaCl; 4, 20 mM MES pH 6.5, 150 mM NaCl; 5, 20 mM bis-tris propane pH 7.0, 150 mM NaCl; 6, 20 mM tris pH 7.5, 150 mM NaCl; 7, 20 mM Tris pH 8.0. **C**) Native PAGE comparative analysis of T4GALA suggests tris results in disassembly while bis-tris propane results in assembly. M, marker; 1, 20 mM Tris pH 7.5, 150 mM NaCl; 2, ; 2, 20 mM bis-tris propane pH 7.5, 150 mM NaCl; 3, 20 mM bis-tris propane pH 7.5, 150 mM NaCl, 5 mM imidazole; 4, 20 mM bis-tris propane pH 7.5, 150 mM NaCl, 25 mM imidazole; 5, 20 mM bis-tris propane pH 7.5, 150 mM NaCl, 250 mM imidazole; 6, 20 mM bis-tris propane pH 7.5, 150 mM NaCl, 50 mM $NH_4Cl$.

#### 3.4 T4GALA assembly and disassembly analyses via SEC

Assembled T4GALA was successfully purified via Ni-NTA purification as described above. Early trial SEC purification attempts indicated the need to purify the target nanocage expediently and to store it in a buffer well-suited for maintaining assembly. Initially, attempts were made to purify via Ni-NTA and then buffer exchanged the target back into the Resuspension Buffer to remove the high imidazole content. By pre-equilibrating the SEC column with Elution Buffer, however, it was determined that the high imidazole ionic strength actually encouraged assembly (**Figure S3**).

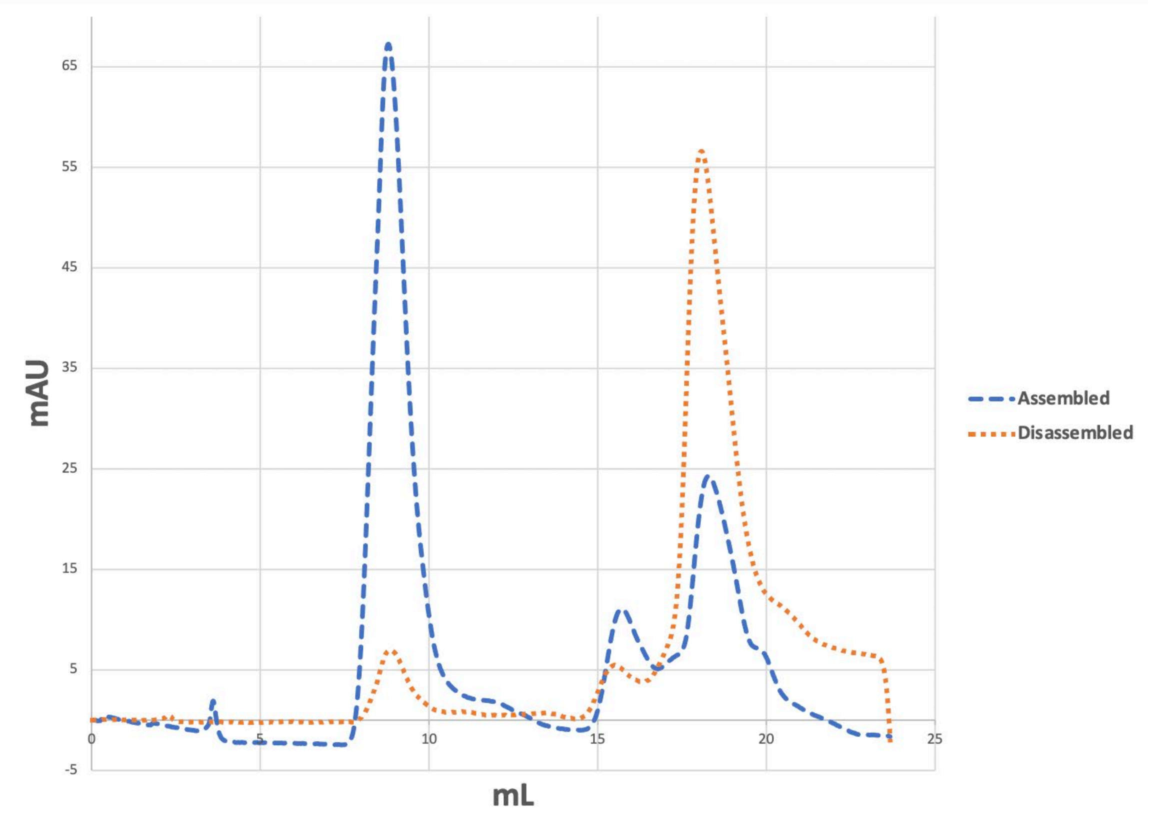

**Figure S3.** Size exclusion chromatography (SEC) analyses of assembled and disassembled T4GALA. The target protein nanocage shifted to a disassembled state when stored overnight in low imidazole-containing Resuspension Buffer (yellow dash). Conversely, by immediately loading post-Ni-NTA purified T4GALA onto an SEC column pre-equilibrated with a relatively higher imidazole buffer (Elution Buffer), a substantially higher portion of the sample remained assembled (blue dash).

### 3.5 Transmission electron microscopy analysis of (cargo-loaded) T4GALA

Various T4GALA and T4GALA-encapsulated constructs were imaged on multiple occasions via TEM to verify assembly of T4GALA throughout the different experiments.

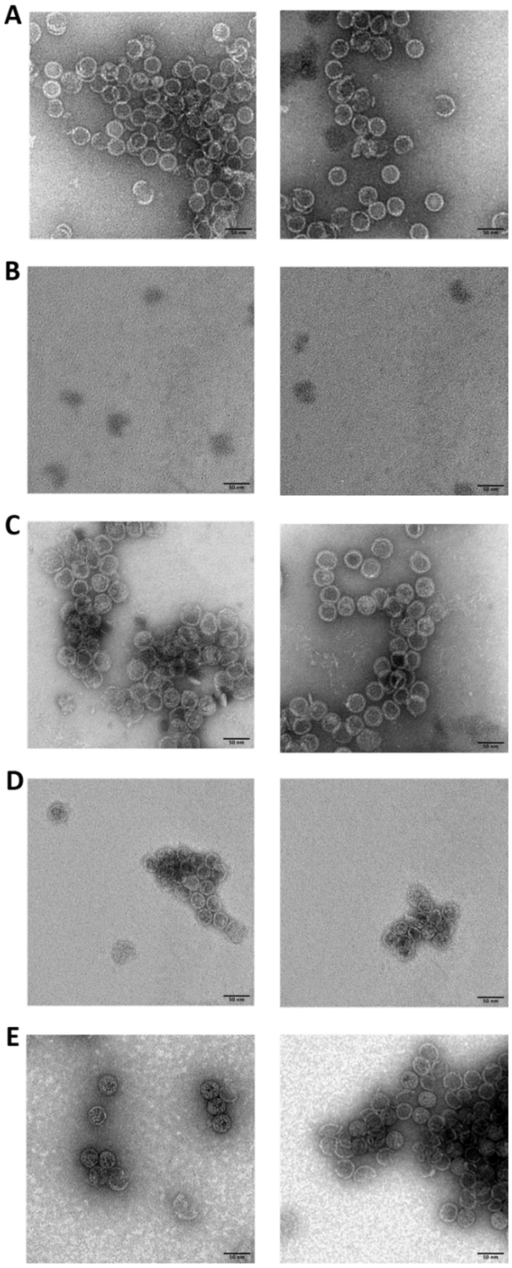

**Figure S4.** TEM analysis of T4GALA and T4GALA-encapsulated constructs. A) Additional representative TEM micrographs of assembled T4GALA. B) Additional representative TEM micrographs of disassembled T4GALA. Notable accumulation of what may be disassembled protomers is observable. C) Additional representative TEM micrographs of reassembled T4GALA. D) Representative TEM micrographs of assembled FADGDH_GALA. E) Representative TEM micrographs of assembled GFP11_T4GALA. It appears a notable tendency exists for T4GALA to aggregate more than QtEnc, possibly due to the hydrophobic nature of the GALA insert. Black scale bars represent 50 nm.

#### 3.6 Cryogenic microscopy analysis of T4GALA

Cryo-EM analysis of T4GALA showed an absence of density corresponding to the engineered GALA insert trigger at the E-loop along the three-fold axis (**Figure S5 and Figure S6**).

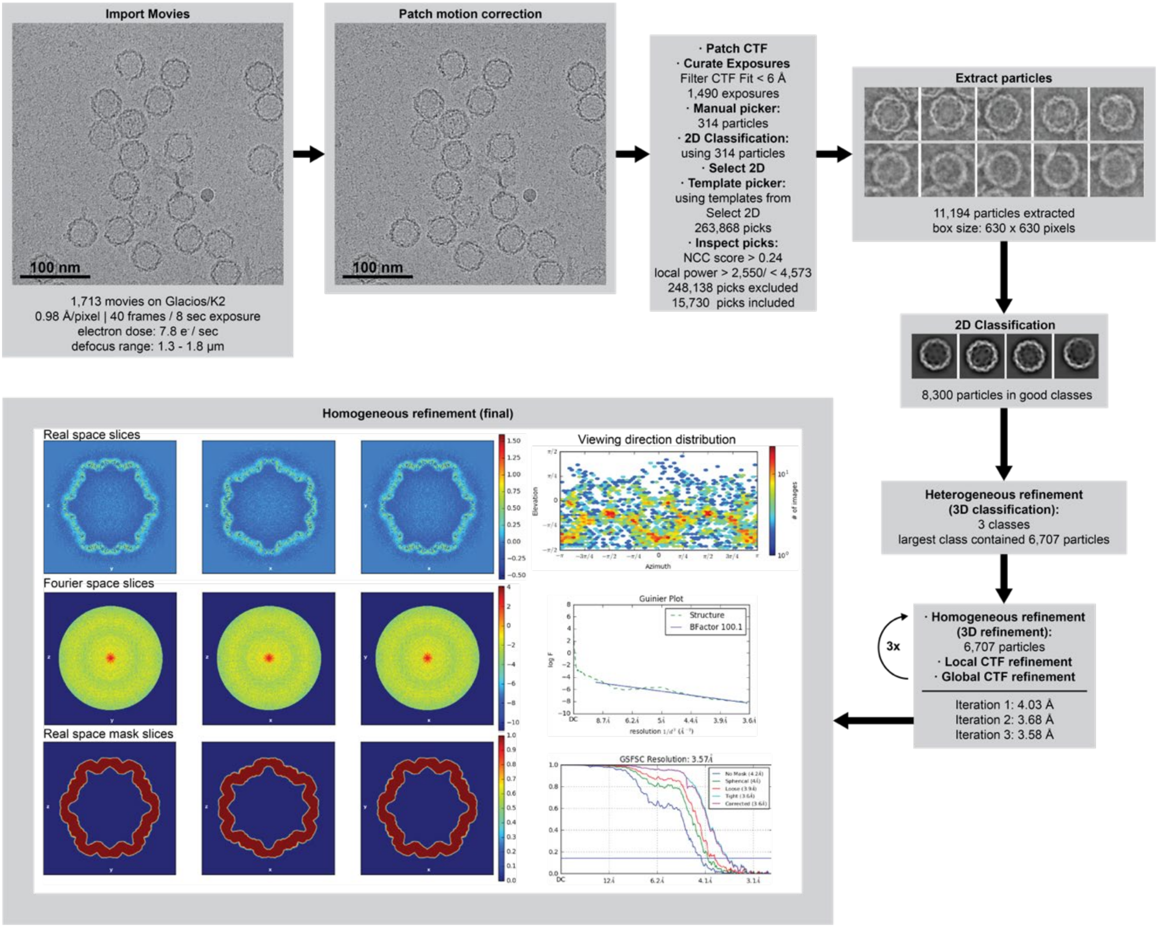

**Figure S5.** Cryo-EM processing pipeline.

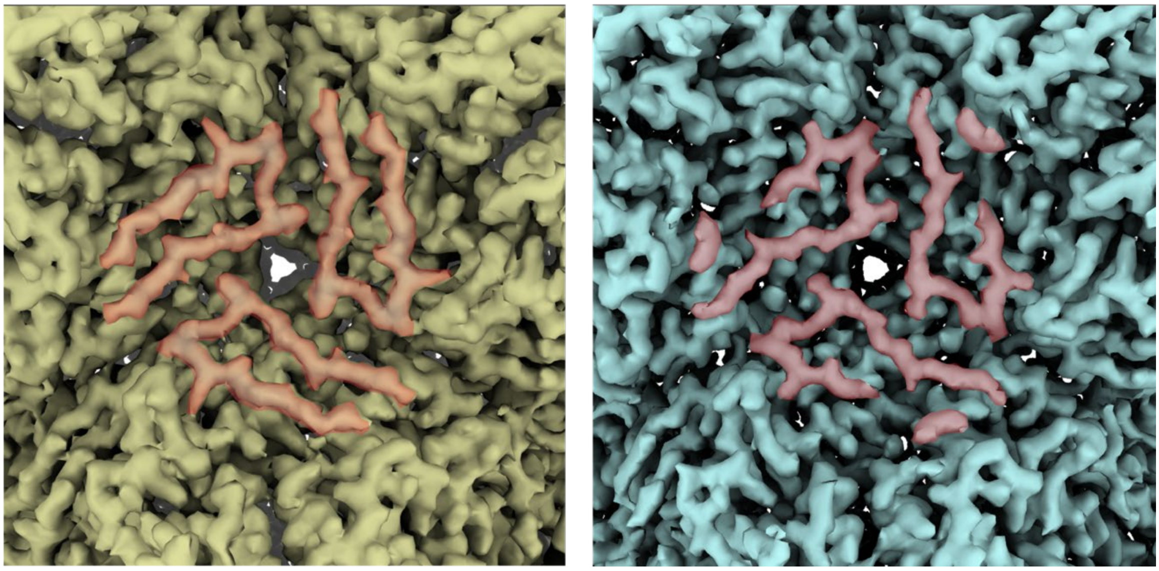

**Figure S6.** Cryo-EM density comparison of QtEnc (PDB 6NJ8; yellow, left) with T4GALA (blue, right) along the three-fold axis showcasing apparent differences in E-loop density (red) and highlighting a lack of density for the T4GALA trigger insert.

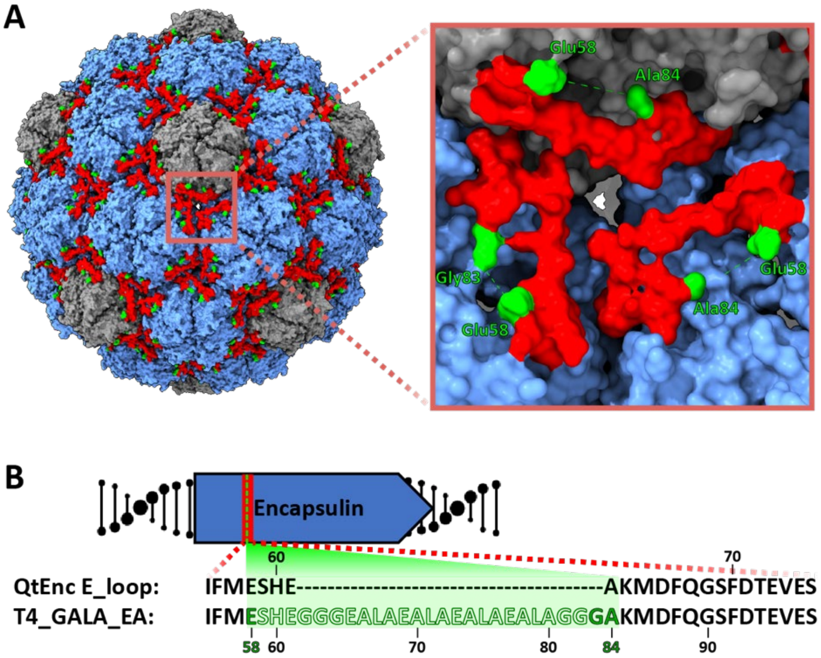

**Figure S7.** Cryo-EM analysis of T4GALA along the pseudo-three-fold axis formed by two neighboring hexameric and one pentameric capsomers. A) Surface diagram of the assembled T4GALA engineered protein cage. Hexameric and petameric facets depicted (blue and grey, respectively) with all E-loops (red) and residues flanking the GALA insert (green) highlighted. Inset (right) depicting the pseudo-three-fold axis to emphasize E-loops (red), including insert-flanking residues Glu58, Gly83, and Ala84 (originally Ala62 from PDB 6NJ8; green) and unresolved E-loop and GALA insert residues 59-82 (green dash). B) Schematic diagram of encapsulin core operon showing the E-loop (red) and highlighting the residues with missing cryo-EM density (silhouettes) between residues with density, Glu58, Gly83, and Ala84 (originally Ala62 from PDB 6NJ8; green).

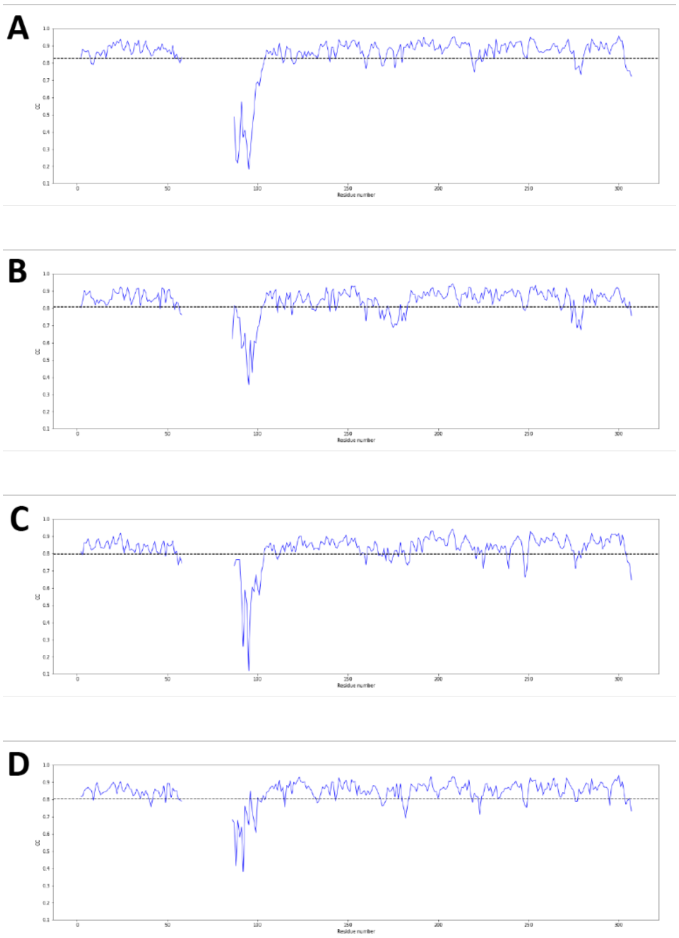

**Figure S8.** Model-to-map correlation analysis of the E-loop arms. Low model-to-map correlation corroborating the dynamic nature of the designed GALA trigger region within the T4GALA E-loop. Panels A through D represent each respective chain.

**Table S2.** Cryo-EM data collection and model building statistics.

| | |
|---|---|
| **Data collection and processing** | |
| Electron microscope | Glacios |
| Voltage (kV) | 200 |
| Total Dose ($e^-$/Å$^2$) | 62 |
| Defocus range (um) | -1.3 to -1.8 |
| Voltage (kV) | 200 |
| Particles | 6707 |
| Resolution (Å) | 3.57 |
| Map sharpening B-factor (Å$^2$) | -100.1 |
| Symmetry for map | I |
| Number of Movies | 1259 |
| **Structure Refinement (ASU)** | |
| Structure B factor (mean) | 81.22 |
| Mean CC for Protein (Mask) | 0.82 |
| Chains in ASU | 4 |

| | |
|---|---|
| Bond lengths (Å) | 0.004 |
| Bond Angles (°) | 0.899 |
| **Ramachandran Statistics** | |
| Favored % | 95.63 |
| Allowed % | 4.37 |
| Disallowed % | 0.00 |
| Rotamer Outliers (%) | 4.3 |
| **MolProbity Score** | 1.95 |
| **Clash Score** | 3.78 |

### In vitro cargo loading analysis

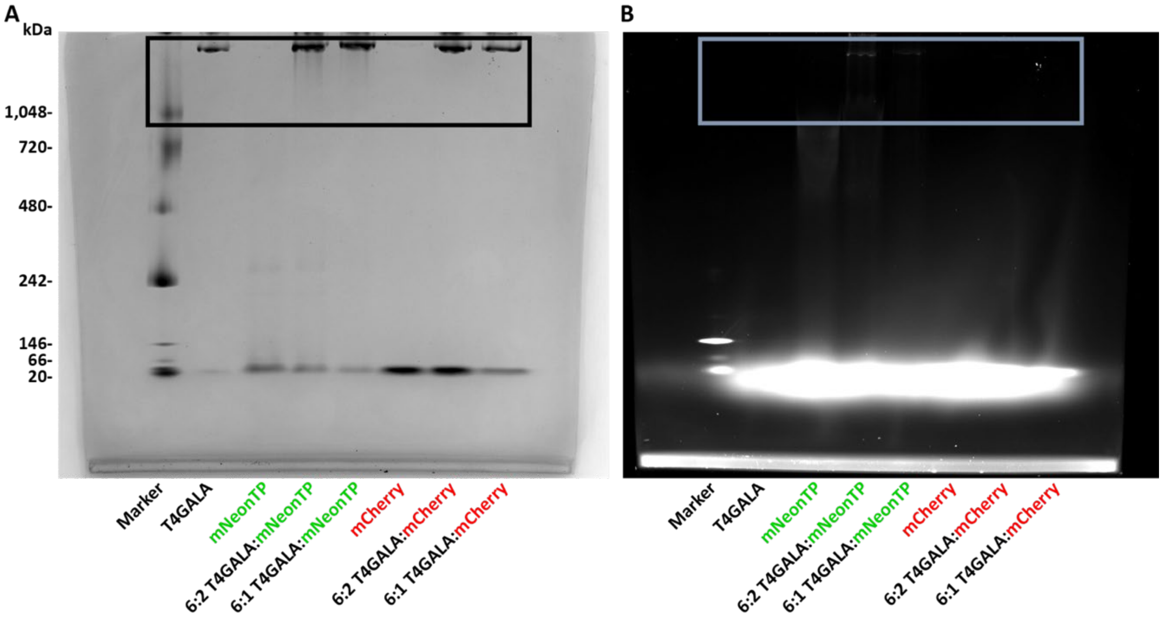

**Figure S9.** *In vitro* cargo loading of T4GALA. A) Full Native PAGE analysis showing high assembled encapsulin bands via Coomassie blue staining (black box). B) Corresponding fluorescence detected along with assembled T4GALA protein band due to targeting peptide-fused mNeon cargo samples. Additionally, a lack of fluorescence is observed from non-encapsulated mCherry treated in parallel. M, marker; T4GALA, T4GALA engineered protein cage; mNeonTP, T4 targeting peptide-fused mNeon fluorescent protein; mCherry, mCherry fluorescent protein without targeting peptide fusion.

### 5. Author Contributions

A.S.C., J.A.J., and T.W.G. designed the project. A.S.C. and T.W.G. designed the engineered protein cage. A.S.C. and J.A.J. conducted the laboratory experiments and transmission electron microscopy, while M.P.A. obtained and analyzed cryo-electron microscopy data. J.A.J. and T.W.G. wrote the manuscript. T.W.G. oversaw the project in its entirety.
